# Development of zygotic and germline gene drives in mice

**DOI:** 10.1101/2020.06.21.162594

**Authors:** Chandran Pfitzner, James Hughes, Melissa White, Michaela Scherer, Sandra Piltz, Paul Thomas

**Affiliations:** School of Biological Sciences, The University of Adelaide, Adelaide, South Australia, Australia, 5005; Precision Medicine, South Australian Health and Medical Research Institute, Adelaide, South Australia, Australia, 5000; School of Medicine, The University of Adelaide, Adelaide, South Australia, Australia, 5005; Robinson Research Institute, The University of Adelaide, Adelaide, South Australia, Australia, 5005

## Abstract

CRISPR-based synthetic gene drives have the potential to deliver a more effective and humane method of invasive vertebrate pest control than current strategies. Relatively efficient CRISPR gene drives have been developed in insects and yeast but not in mammals. Here we investigated the efficiency of CRISPR/Cas9-based gene drives in *Mus musculus* by constructing “split drive” systems with Cas9 under the control of zygotic (CAG) or germline (Vasa) promoters. While both systems generated double stranded breaks at their intended target site *in vivo*, no homing was detectable. Our data indicate that robust and specific Cas9 expression during meiosis is a critical requirement for the generation of efficient CRISPR-based synthetic gene drives in rodents.

## Introduction

Invasive rodents, including mice, pose a significant threat to biodiversity, particularly on islands, and are the likely cause of many species extinctions (Blackburn *et al*. 2004; Harris 2008; Doherty *et al*. 2016). The economic burden to the agriculture industry is also considerable, costing tens of millions of dollars to many countries each year (Stenseth *et al*. 2003). Previous attempts at invasive vertebrate pest control have had some success, but there are still many challenges including cost and ethical considerations (Gregory *et al*. 2014).

Manipulation of natural gene drives has been proposed as a tool for population suppression of invasive rodent pests by rapidly spreading a gene through a wild population that would ultimately have a deleterious effect on reproductive fitness. These include transposable elements (TEs), homing endonuclease genes (HEGs) and meiotic drives. However, all these have significant drawbacks; TEs copy themselves to unpredictable locations causing their own repression or disruption of endogenous genes (Sinkins and Gould 2006), HEGs are not found in animals and rely on a site-specific nuclease that can’t be tailored to a specific location (Burt and Koufopanou 2004) and meiotic drives are rare, poorly understood and generally very large, making them difficult to manipulate (Kelemen and Vicoso 2018; Charron *et al*. 2019).

Synthetic, clustered regularly interspaced short palindromic repeats (CRISPR)-based gene drives provide an alternative that has a small genomic footprint, can be inserted almost anywhere in the genome and can be created relatively quickly and easily. CRISPR gene drives are composed of a cassette integrated into a specific genomic site that expresses CRISPR-associated protein 9 (Cas9) endonuclease and a customisable guide RNA (gRNA) designed to cut the homologous WT locus. Repair of the double stranded break (DSB) by homology directed repair (using the gene drive allele as a repair template) results in conversion of the WT allele to a gene drive allele, in a process termed “homing”, which renders the cell homozygous for the gene drive allele. Homing can be restricted to the gamete precursors resulting in selective homozygosity in the germline whilst the somatic cells remain heterozygous. The homing event will ensure that the gene drive allele will be present in all of the gametes and will be transmitted to all of that organism’s progeny. Thus, over several generations, gene drives will rapidly spread through a given population. Alternatively, repair of the DSB by error-prone pathways such as non-homologous end joining, microhomology-mediated end joining and single-strand annealing can generate an indel that interrupts the gRNA binding sequence thereby creating a mutant allele that is immune to homing. These so called “resistant” alleles pose a significant barrier to gene drive spread (Prowse *et al*. 2017).

Highly efficient CRISPR gene drives have already been generated in the following species, where germline homing rates (in brackets) are defined as the percentage of WT alleles converted to gene drive alleles: *Drosophila melanogaster* (>76%) (Gantz and Bier 2015; Champer *et al*. 2017), *Anopheles stephensi* (>96%) (Gantz *et al*. 2015), *Saccharomyces cerevisiae* (>99%) (DiCarlo *et al*. 2015) and *Anopheles gambiae* (>82%) (Hammond *et al*. 2016). While gene drives are yet to be developed in mice, Grunwald *et al*. (2019) recently generated a “single generation” homing system. Highly variable and relatively inefficient (0-72%) homing was observed in females but no homing was detected in males. While this study suggests that it may be possible to develop deployable gene drives in mice, key components including the choice of promoter for Cas9 expression are yet to be assessed.

Here, we describe the first attempt to develop mice with functional, multi-generational gene drives under the control of either zygotic or germline promoters. To ensure that the mice we generated did not pose any threat to the environment if unintentionally released (Akbari *et al*. 2015), we employed a “synthetic target” strategy as a molecular safeguard. In addition, we used a “split drive” system where Cas9 is completely separated from the gRNA-containing homing cassette.

## Materials and Methods

### Mouse Model Generation

*Cas9* mRNA for zygotic injections was generated from the XhoI-digested pCMV/T7-hCas9 plasmid (Toolgen) using the mMESSAGE mMACHINE® T7 ULTRA Transcription Kit (Ambion) and purified using RNeasy Mini Kit (Qiagen).

The Tyr-gRNA for zygotic injection of constructs targeting Tyrosinase (*Tyr*) intron 1 was designed using the Zhang lab tool (Hsu *et al*. 2013). The Rosa26-gRNA design was previously published (Platt *et al*. 2014). gRNA-containing plasmids were generated in pSpCas9(BB)-2A-Puro (PX459) V2.0 (Addgene; 62988) (Ran *et al*. 2013) with guide oligonucleotides purchased from Sigma/IDT. gRNA was generated with T7 promoter-containing oligos (Sigma/IDT) using HiScribe™ T7 Quick High Yield RNA Synthesis Kit (NEB) and purified using RNeasy Mini Kit (Qiagen).

The Tyr^gRNA-Tomato^ plasmid was purchased from GenScript and prepared using PureLink™ HiPure Plasmid Midiprep Kit (Invitrogen). Plasmid (10 ng/µL), *Cas9* mRNA (25 ng/µL), and Tyr-gRNA (10 ng/µL) were injected into the pronuclei of C57BL/6J zygotes, transferred to pseudo-pregnant recipients, and allowed to develop to term.

An unanticipated side effect of the presence of the *Tyr*^*gRNA-Tomato*^ allele resulted in the sudden and largely unexplained deaths of mice carrying that allele. Due to the ethical concerns surrounding this unexpected toxicity, the *Tyr*^*gRNA-Lite*^ mouse was designed and generated, containing only the U6-Neo-gRNA cassette from the *Tyr*^*gRNA-Tomato*^ mouse.

Tyr^gRNA-Lite^ ssDNA was purchased from IDT as a Megamer®. ssDNA (10 ng/µL), PNA Bio Cas9 protein (50 ng/µL) and Tyr-gRNA (25 ng/µL) were injected into the pronuclei of C57BL/6J zygotes, transferred to pseudo-pregnant recipients, and allowed to develop to term.

Tyr^Target^ dsDNA was purchased from IDT. dsDNA (100 ng/µL), c*as9* mRNA (12.5 ng/µL) and Tyr-gRNA (5 ng/µL) were injected into the cytoplasm of C57BL/6J zygotes, transferred to pseudo-pregnant recipients, and allowed to develop to term.

*Gt(ROSA)26Sor*^*tm1*.*1(CAG-cas9*,-EGFP)Fezh*^/J mice (*Rosa26*^*Cas9*^ mice) were supplied by JAX (Platt *et* al. 2014).

Vasa-Cas9 dsDNA was constructed as follows. Gibson assembly was used to clone the Vasa-β-globin-II fragment from the Vasa-Cre plasmid (Addgene; 15885) (Gallardo *et al*. 2007) and the Cas9-BGH fragment from pSpCas9(BB)-2A-Puro (PX459) V2.0 (Addgene; 62988) (Ran *et* al. 2013) into the pStart-K plasmid (Addgene; 20346) (Wu *et al*. 2008). This plasmid was expanded with PureLink™ HiPure Plasmid Midiprep Kit (Invitrogen), digested with BamHI (NEB), purified using Gel DNA Recovery Kit (Zymoclean), and further purified on a floating dialysis membrane for 2.5 h. dsDNA (3 ng/µL), PNA Bio Cas9 protein (50 ng/µL) and Rosa26-gRNA (25 ng/µL) were injected into the pronuclei of C57BL/6J zygotes, transferred to pseudo-pregnant recipients, and allowed to develop to term.

### Mouse crosses

*Rosa26*^*Cas9*^/+ ; *Tyr*^*gRNA-Tomato*/*Target*^ mice were generated by crossing *Rosa26*^*Cas9/Cas9*^ ; *Tyr*^*Target/Target*^ mice to *Tyr*^*gRNA-Tomato*^/+ mice and also by crossing *Rosa26*^*Cas9/Cas9*^ ; *Tyr*^*gRNA-*^ _*Tomato*_/+ mice to *Tyr*^*Target/Target*^ mice.

*Vasa-Cas9-2*/+ ; *Tyr*^*gRNA-Tomato*/*Target*^ mice were generated by crossing *Vasa-Cas9-2*/*Vasa-Cas9-2* ; *Tyr*^*Target*/*Target*^ mice to *Tyr*^*gRNA-Tomato*^/+ mice. *Vasa-Cas9-4*/+ ; *Tyr*^*gRNA-Lite*/*Target*^ mice were generated by crossing *Vasa-Cas9-4*/+ ; *Tyr*^*Target*/*Target*^ mice to *Tyr*^*gRNA-Lite/gRNA-Lite*^ mice.

Sequencing of *Tyr*^*Target*^ loci in *Vasa-Cas9-2*/+ ; *Tyr*^*gRNA-Tomato*/*Target*^ and *Vasa-Cas9-4*/+ ; *Tyr*^*gRNA-Lite*/*Target*^ mice confirmed no carryover of maternal *Cas9* mRNA into eggs as no indels were seen.

### DNA extractions

gDNA was extracted from embryo, tail tip, or ear notch biopsies using the High Pure PCR Template Preparation Kit (Roche), KAPA Express Extract kit (Roche), or MyTaq™ Extract-PCR Kit (Bioline). gDNA was prepared from blastocysts in a 20 µL solution of 185.5 mM pH 8.3 Tris-HCl, 185.5 mM KCl, 7.4×10^−6^ % gelatin, 8.3×10^−4^ % Polysorbate 20, 1.48 % tRNA from baker’s yeast (Sigma), 1.15 mg/mL Proteinase K (Thermo Scientific) which was incubated at 56 °C for 10 min and 95 °C for 10 min.

gDNA was prepared from sperm by washing epididymides in PBS at 37 °C followed by transfer to the centre well of a 37 °C Center-Well Organ Culture Dish (Falcon) containing 500 µL of M2 medium (Sigma) in the centre well and 3 mL of PBS in the outer well. Several incisions were made across epididymides before incubating at 37 °C/5% CO_2_ for 5 min. The sperm-containing M2 medium was then centrifuged at 400 RCF for 10 min. The supernatant was discarded, 500 µL PBS was added and then centrifuged at 8000 RCF for 1 min. The supernatant was discarded and the pellet resuspended in 100 µL PBS. 400 µL Tissue Lysis Buffer (Roche) and 50 µL Proteinase K (Roche) was added and the solution was vortexed. The solution was then incubated at 55 °C for 1 h, 50 µL of 1 M DTT was added, vortexed rigorously, then incubated O/N at 55 °C. High Pure PCR Template Preparation Kit (Roche) was then used for DNA extraction, eluting in 80 µL EB.

### RNA Extraction

Acid guanidinium thiocyanate-phenol-chloroform RNA extraction was performed on testes, ovaries, and spleens. RNA was purified using the RNeasy Mini/Micro kit (Qiagen) in conjunction with RNase-Free DNase Set (Qiagen). cDNA was generated using the High-Capacity RNA-to-cDNA™ Kit (Applied Biosystems).

### Genotyping Analysis

PCRs were performed using either Taq DNA Polymerase (Roche), Phusion HF DNA Polymerase (NEB) or KAPA2G Fast Genotyping Mix (Roche) using their associated buffers or alternatively FailSafe PCR PreMix buffers (Epicentre). Fast SYBR Green Master Mix (Applied Biosystems) was used for qPCRs. Primers were purchased from IDT and Sigma. All primers used are shown in **Supplemental Table 3**.

50 ng genomic DNA was digested with MseI for droplet digital PCR (ddPCR). ddPCR was performed with 12.5 ng digested DNA, 1.25 µL of control assay targeting Rpp30 (Bio-Rad), 1.25 µL of target assay targeting U6-Neo-gRNA (Bio-Rad) and 12.5 µL ddPCR Supermix for Probes (No dUTP) (Bio-Rad). Droplets were generated with QX200 Droplet Generator (Bio-Rad) and read with QX200 Droplet Reader (Bio-Rad).

Sanger sequencing was performed by Australian Genome Research Facility (Adelaide, Australia).

Indels at *Tyr*^*Target*^ were detected by RFLP in combination with T7 Endonuclease digestion to cleave heteroduplexes.

## Results

### Design and generation of the homing system

To generate a “synthetic target” site, we identified a candidate homing gRNA (Neo-gRNA) targeting the bacterial kanamycin kinase gene and confirmed its on-target cleavage activity using embryonic stem cells harbouring that gene (**Supplemental Figure 1**). Next, we generated a “donor” mouse line (*Tyr*^*gRNA-Tomato*^) by inserting a Neo-gRNA expression cassette (Cong *et al*. 2013) and a ubiquitous dTomato reporter gene (Shaner *et al*. 2004) into the first intron of the *Tyr* gene (**Figure 1A**, **Supplemental Figure 2A**,**B**). To generate a null allele, we included a 3x SV40 polyadenylation signal in the same orientation as *Tyr* so that the *Tyr* transcript is subject to premature termination. Inactivation of *Tyr* was confirmed by the white coat of *Tyr*^*gRNA-Tomato*^ homozygotes (**Supplemental Figure 2C**). We also generated a complementary “receiver” mouse line (*Tyr*^*Target*^) with the Neo-gRNA target sequence at the same locus as *Tyr*^*gRNA-Tomato*^ (**Figure 1C**). We reasoned that by targeting an intronic location, indel mutations generated by error-prone repair pathways would not significantly reduce tyrosinase (TYR) function. Therefore, conversion of the *Tyr*^*Target*^ allele to a *Tyr*^*gRNA-Tomato*^ allele via zygotic homing would generate a white mouse whereas error-prone repair pathways would generate a black mouse. As expected, the *Tyr*^*Target*^ insertion did not detectably alter TYR activity as demonstrated by the black coat of *Tyr*^*Target*^ homozygous mice (**Supplemental Figure 3**).

**Figure 1.**
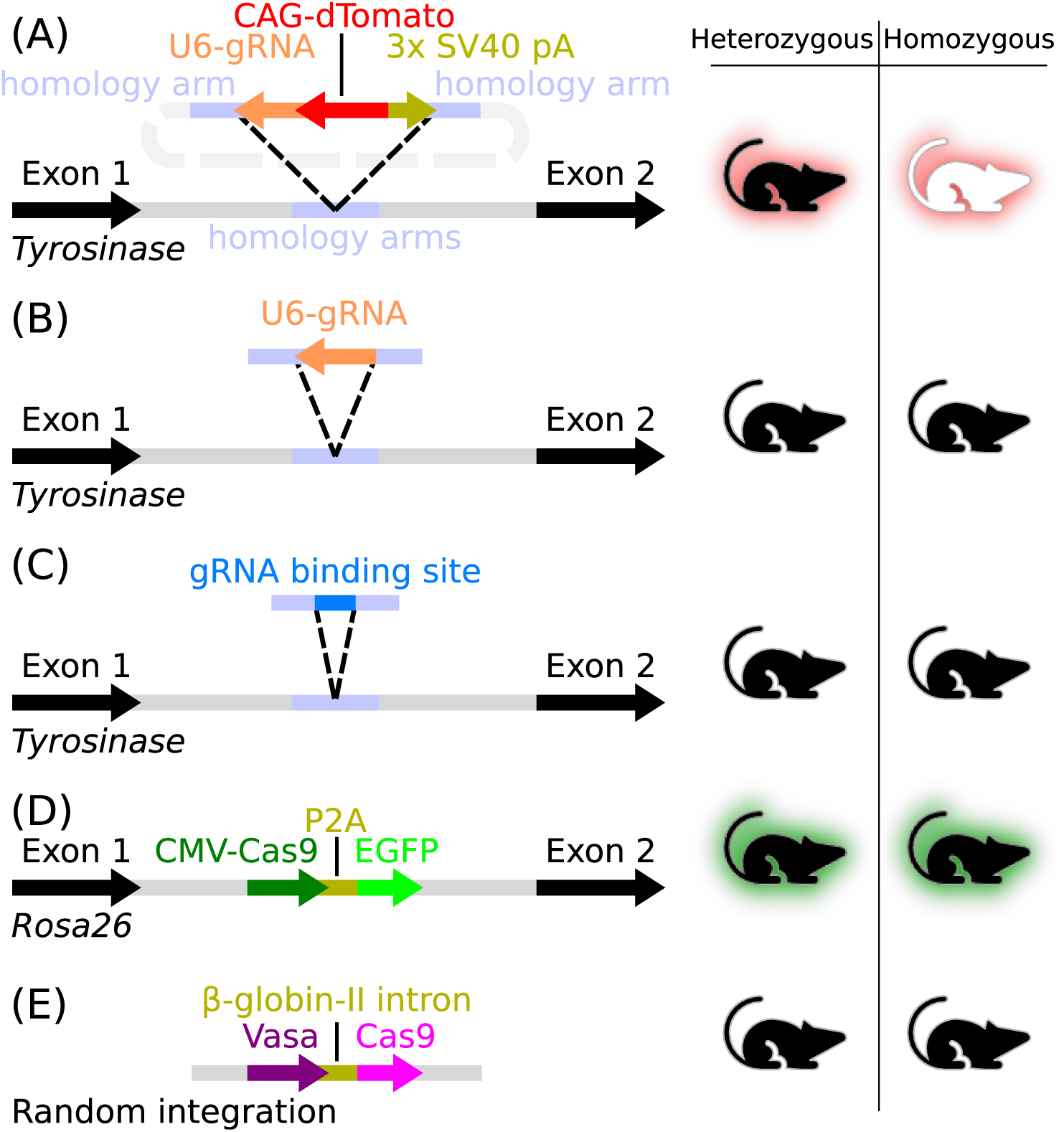
Mouse models. **(A)** *Tyr*^*gRNA-Tomato*^ mice contained a U6-driven Neo-gRNA, a CAG-driven dTomato gene and a 3x SV40 polyA signal inserted into *Tyr* intron 1. **(B)** *Tyr*^*gRNA-Lite*^ mice contained a U6-driven Neo-gRNA inserted into the same location as (A) in *Tyr* intron 1. **(C)** *Tyr*^*Target*^ mice contained the Neo-gRNA target sequence and NGG PAM inserted into the same location as (A) in *Tyr* intron 1. **(D)** The *Rosa26*^*Cas9*^ mouse contained a CAG-driven *cas9* linked via P2A to *EGFP* in the Rosa26 locus (Platt *et al*. 2014). **(E)** The *Vasa-Cas9* mouse lines contained a Vasa-driven *hSpCas9* linked via the β-globin-II intron (non-targeted integration). Coat colour and fluorescence phenotypes are shown on the right.

**Figure 2.**
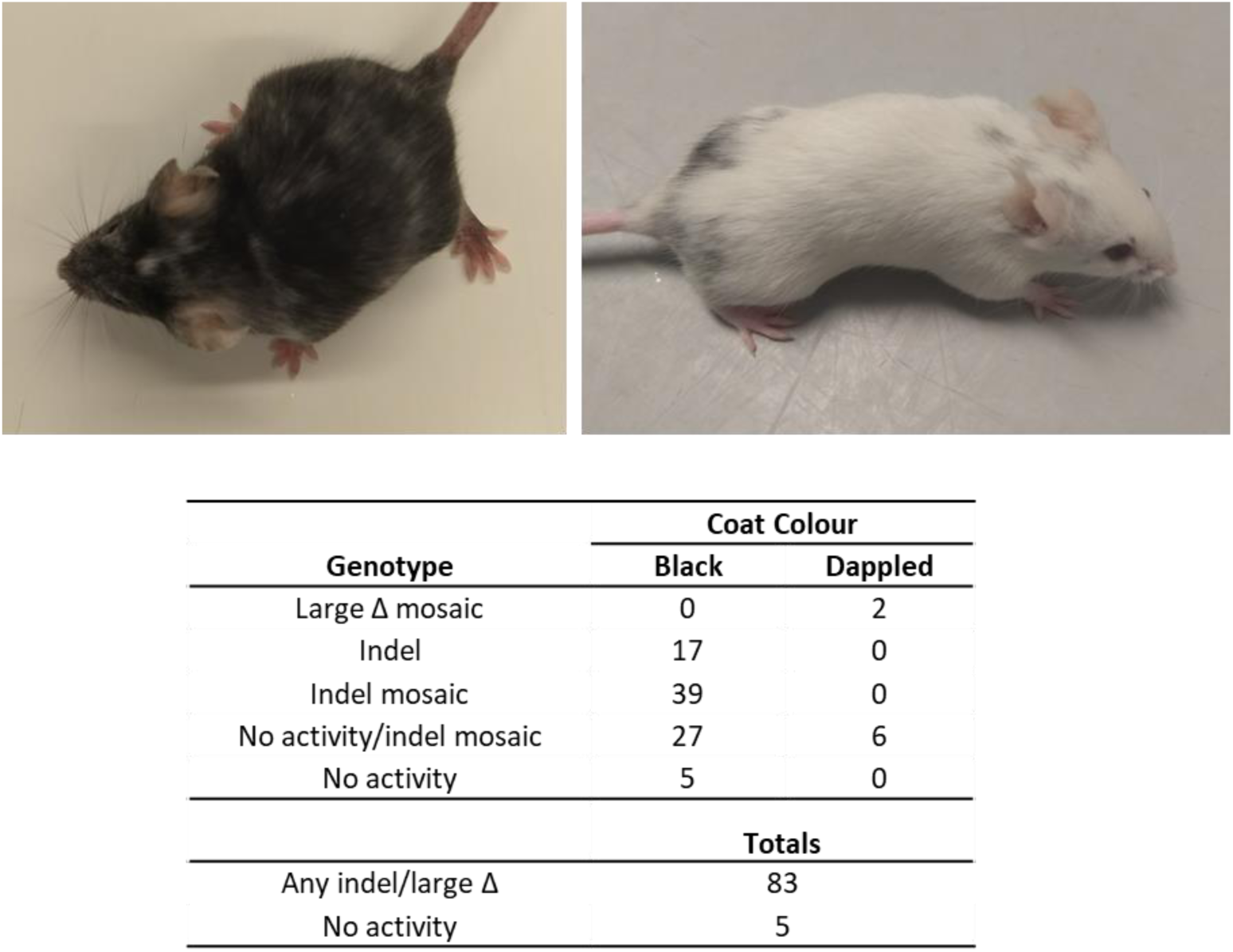
Zygotic-homing gene drive results. **Top:** Representative *Rosa26*^*Cas9*^/+ ; *Tyr*^*gRNA-Tomato/Target*^ mice with dappled coats showing wide variability in mosaicism. **Bottom:** Table of genotypes of *Tyr*^*Target*^ allele(s) in *Rosa26*^*Cas9*^/+ ; *Tyr*^*gRNA-Tomato*/*Target*^ gene drive mice. “Large Δ mosaic” contain a large deletion and 1 or more indels. “Indel” contain a single indel. “Indel mosaic” contain 1 or more indels. “No activity/indel mosaic” contain an uncut *Tyr*^*Target*^ allele and 1 or more indels/large deletions. “No activity” contain a single uncut *Tyr*^*Target*^ allele. Dappled is a mix of black and white fur.

**Figure 3.**
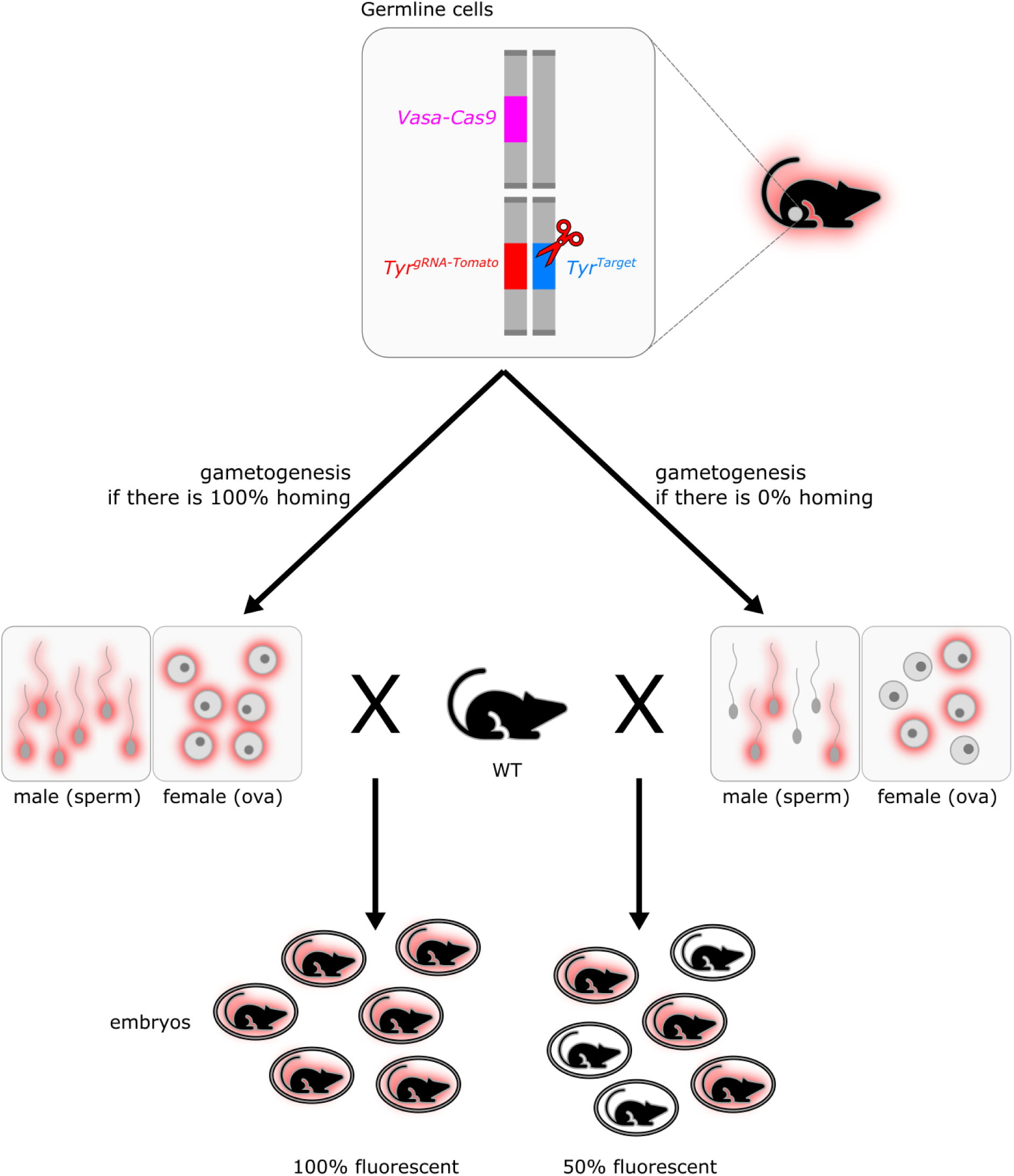
Germline-homing gene drive activity. Experimental mice (top right) contain *Vasa-Cas9, Tyr*^*gRNA-Tomato*^, and *Tyr*^*Target*^ alleles. Somatic tissue will not express Cas9 and thus maintain an intact *Tyr*^*Target*^. Cas9 produced in germline tissue (top) will complex with Neo-gRNA and generate a DSB. DNA repair mechanisms would then either copy the *Tyr*^*gRNA-Tomato*^ allele over from the donor chromosome (homing) or create an indel (*Tyr*^*TargetΔ*^). The sperm and ova produced (middle) are shown in the two most extreme alternative possibilities, 100% or 0% homing. The actual homing percent is calculated after crossing the original mouse with a WT mouse and counting the number of fluorescent offspring.

To assess homing, Cas9 was expressed using a separate transgene. For zygotic homing we used *Rosa26*^*Cas9*^ mice (Platt *et al*. 2014) which express Cas9 (and EGFP) ubiquitously (**Figure 1D**). For germline homing experiments we generated *Vasa-Cas9* lines via random integration (**Figure 1E**).

### Zygotic homing

To assess homing in the zygote, we generated 96 *Rosa26*^*Cas9*^/+ ; *Tyr*^*gRNA-Tomato*/*Target*^ mice. Two groups were identified based on coat colour: black mice (88) and dappled mice (8). 94% (83/88) of the black mice carried indels in the *Tyr*^*Target*^ allele indicating efficient production and high cleavage activity of the Neo-gRNA/Cas9 complex (**Figure 2**). To assess whether homing had occurred in some cells of the dappled mice, we quantitated the *Tyr*^*gRNA-Tomato*^ allele using ddPCR (**Supplemental Figure 4**). Surprisingly, only a single copy of the *Tyr*^*gRNA-Tomato*^ allele was present, indicating a lack of homing. Based on our previous observation that DSB repair in zygotes often generates large (>100 bp) deletions (Adikusuma *et al*. 2018), we amplified the target locus using primers distant from the cleavage site. Large deletions were identified in all dappled mice (**Supplemental Figure 5**). As *Tyr* exon 1 was 500 bp upstream of the target site it seems likely that these deletions extended into exon 1 to generate null alleles consistent with the partial (mosaic) albino phenotype.

**Figure 4.**
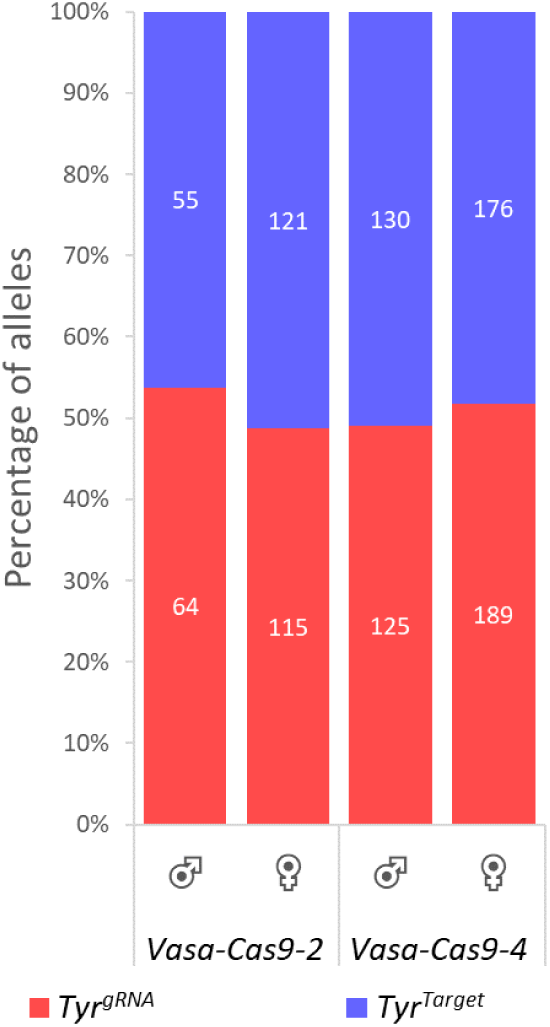
Germline-homing genotyping data. Bar graph showing the percentage of different alleles in the offspring of gene drive mice *Vasa-Cas9-2*/+ ; *Tyr*^*gRNA-Tomato*/*Target*^ and *Vasa-Cas9-4*/+ ; *Tyr*^*gRNA-Lite*/*Target*^. Total number of offspring shown in columns.

**Figure 5.**
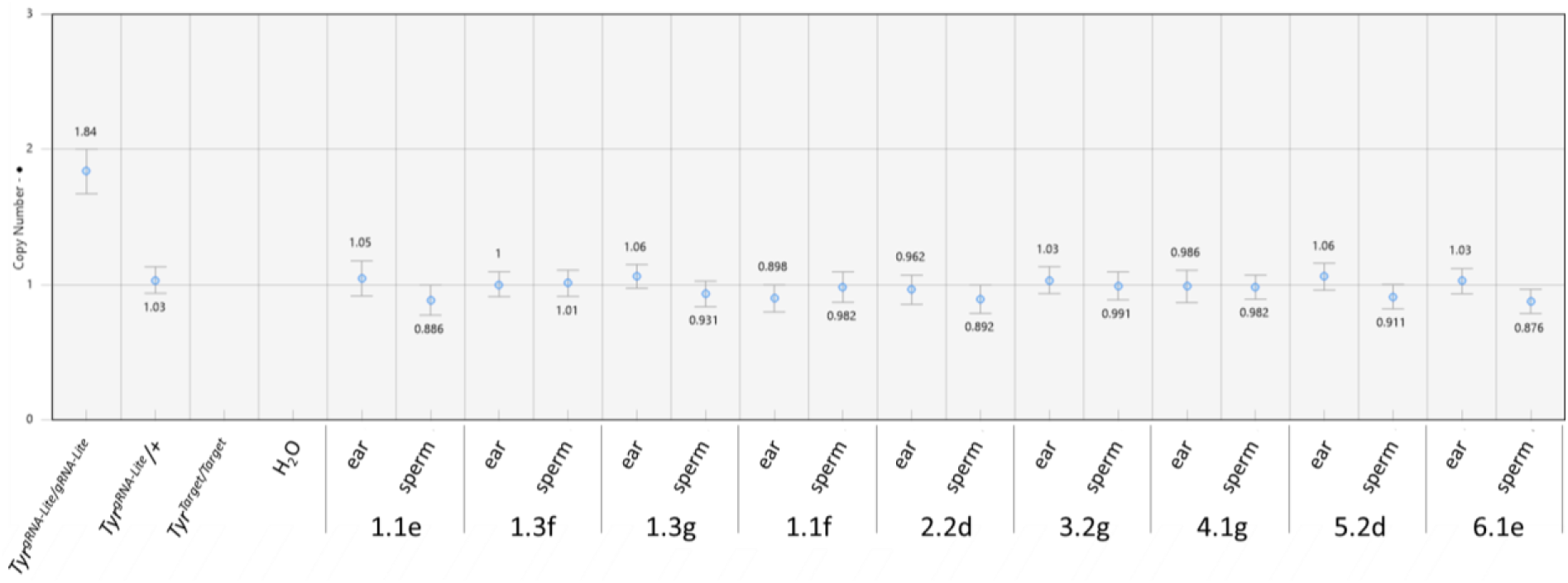
*Tyr*^*gRNA-Lite*^ ddPCR assay. Comparison of ***Tyr***^***gRNA-Lite***^ levels in genomic DNA from somatic tissue (ear) and sperm from *Vasa-Cas9-4/+* ; *Tyr*^*gRNA-Lite*/*Target*^ mice.

Notably, multiple *Tyr*^*Target*^ indel alleles were also detected in many black mice (**Figure 2**) indicating mosaicism. Taken together, these data indicate that despite efficient cleavage of the target sequence, homing did not occur in zygotes or cleavage-stage embryos.

### Germline homing

For germline homing experiments, we initially generated a *Vasa-Cas9* transgenic mouse using the previously characterised 5.6 kb Vasa promoter fragment (*Vasa-Cas9-2*; **Figure 1E**) which has been shown to drive robust Cre expression in the male and female germline (Gallardo *et al*. 2007). RT-qPCR analysis showed that Cas9 was expressed in both testes and ovaries, although, unexpectedly, expression in the latter was extremely low (**Supplemental Figure 6A)**. To assess germline homing, *Vasa-Cas9-2*/+ ; *Tyr*^*gRNA-Tomato*/*Target*^ mice were generated and mated with WT partners. If homing occurs in the germline, >50% of their progeny will carry the *Tyr*^*gRNA-Tomato*^ red fluorescence marker while no homing would result in ∼50% transmission (**Figure 3**). Altogether we screened 355 offspring from 6 *Vasa-Cas9-2*/+ ; *Tyr*^*gRNA-Tomato*/*Target*^ males (119 offspring) and 12 *Vasa-Cas9-2*/+ ; *Tyr*^*gRNA-Tomato*/*Target*^ females (236 offspring; **Figure 4, Supplemental Table 1**). No significant increase (χ^2^ test) in *Tyr*^*gRNA-Tomato*^ transmission was observed in males (53.8%, p=0.46) or females (48.7%, p=0.75). To assess gRNA/Cas9 cleavage activity, non-fluorescent offspring were screened for indels in the *Tyr*^*Target*^ allele by RFLP (**Supplemental Figure 7, Supplemental Table 1**). Only a low percentage of indels were present in progeny from both males (14.5%) and females (9.3%) with the majority of *Tyr*^*Target*^ alleles remaining uncut.

To investigate if higher Cas9 expression levels would promote germline homing, we generated an additional *Vasa-Cas9* mouse line (*Vasa-Cas9-4*). Expression analysis showed a similar Cas9 mRNA level to *Vasa-Cas9-2* in the testes but ∼30 fold higher Cas9 mRNA levels in ovaries (**Supplemental Figure 6A**), possibly due to the higher transgene copy number (**Supplemental Figure 6B**). Germline homing experiments were performed as described for *Vasa-Cas9-2* except with a shorter donor construct (Tyr^gRNA-Lite^) lacking the dTomato expression cassette (see Materials and Methods). Offspring from *Vasa-Cas9-4/+* ; *Tyr*^*gRNA-Lite*/*Target*^ x WT matings were genotyped by PCR (**Figure 4, Supplemental Table 2**). Altogether we screened 620 offspring from 10 transgenic males (255 offspring) and 10 transgenic females (365 offspring). Similar to *Vasa-Cas9-2*, no significant increase (χ^2^ test) in *Tyr*^*gRNA-Lite*^ transmission above 50% was detected from transgenic males (49.0%, p=0.80) or females (51.8%, p=0.53). 87.1% of *Tyr*^*Target*^/+ male offspring carried indels at the target locus compared with 11.0% of *Tyr*^*Target*^/+ female offspring (**Supplemental Figure 7A, Supplemental Table 2**). Finally, to investigate whether a low level of homing was occurring in *Vasa-Cas9-4/+* ; *Tyr*^*gRNA-Lite*/*Target*^ male mice, we investigated whether >50% of sperm carried the *Tyr*^*gRNA-Lite*^ allele using ddPCR (**Figure 5**). No difference in *Tyr*^*gRNA-Lite*^ level between sperm and control somatic (ear) was detected, indicating an absence of homing.

## Discussion

### Zygotic Homing

While the vast majority of gene drive experiments have employed germline promoters to drive Cas9, it is also notionally possible for homing to occur in the zygote. Indeed, targeted integration of dsDNA templates can occur (albeit inefficiently) in zygotes, confirming that HDR is active in the early embryo (Yang *et al*. 2013). Further, it has recently been reported that “interhomolog repair” (effectively the same process as gene drive homing), can occur in human embryos (Ma *et al*. 2017), although concerns have been raised about the interpretation of these results (Egli *et al*. 2018) and (Adikusuma *et al*. 2018). In our experiments using the constitutive CAG promoter, we find no evidence of homing in zygotes despite the high efficiency of DSB generation (>95% of *Rosa26*^*Cas9*^/+ ; *Tyr*^*gRNA-Tomato*/*Target*^ mice have indels). Thus, we conclude that error-prone repair pathways predominate over HDR in the early embryo, consistent with the recent observations of (Grunwald *et al*. 2019). While this DSB repair bias may reflect the availability of endogenous DNA repair proteins, the separation of the donor and receiver alleles into distinct pronuclei until after G_2_ phase (18-20 hours after fertilisation) (Ciemerych and Sicinski 2005) would also limit the opportunity for zygotic homing to occur.

Approximately 77% of *Rosa26*^*Cas9*^/+ ; *Tyr*^*gRNA-Tomato*/*Target*^ mice were mosaic, having multiple *Tyr*^*Target*^ alleles, including many with >3 alleles. Zygotic chromosomes are transcriptionally repressed until the G_2_ phase (Jukam *et al*. 2017) which delay the formation of the gRNA/Cas9 complex until after S phase or cell division, resulting in mosaicism. The strength of the CAG and U6 promoters may also be limiting, resulting in insufficient gRNA/Cas9 complex formation and/or translocation to the *Tyr*^*Target*^ locus in the zygote. It is also possible that flawless NHEJ-mediated DNA repair occurs in a proportion of zygotes thereby delaying the generation of indels until the 2-cell stage or later. Surprisingly, given that the Cas9 and gRNA promoters are constitutive, 5% of *Rosa26*^*Cas9*^/+ ; *Tyr*^*gRNA-Tomato*/*Target*^ mice had an intact *Tyr*^*Target*^ allele and a further 34% had a mix of both intact alleles and indels. This suggest that flawless NHEJ-mediated DNA repair occurs at a relatively high level in somatic cells. In the 5 mice that were free of indels, continued production of EGFP (linked to Cas9 via P2A) indicates that Cas9 protein is still being produced, and continued production of dTomato tells us there is no gene silencing happening in the region of Neo-gRNA transcription (**Supplemental Figure 8**), although it is possible that specific downregulation of the U6 promoter (for Neo-gRNA) may be occurring or increased Cas9 mRNA/protein degradation may occur through unknown mechanisms.

It is possible that carryover of maternal mRNA or protein could have influenced the probability of homing and the developmental stage at which indels are generated. However, we found no evidence for a difference in indel frequency or timing in *Rosa26*^*Cas9*^/+ ; *Tyr*^*gRNA-Tomato*/*Target*^ mice based on the parental origin of the Cas9 or gRNA expression alleles (**Supplemental Figure 9**).

### Germline Homing

Studies in *D. melanogaster, A. stephensi* and *A. gambiae* have shown that homing can occur with high efficiency when Cas9 expression is driven in the germline using promoter fragments (Gantz and Bier 2015; Gantz *et al*. 2015; Hammond *et al*. 2016). Here we used a similar strategy to express Cas9, employing the 5.6kb murine Vasa proximal promoter, which is one of the few characterised mammalian promoter fragments that is expressed in the male and female germline (Gallardo *et al*. 2007). In contrast to insects, transmission of the donor allele was not statistically higher than 50% in either of the Vasa-Cas9 lines. While this does not completely rule out homing, it is clear that it is not occurring at a substantial or useful level.

Although significant homing did not occur, sufficient Cas9 was produced to generate indels in the *Tyr*^*Target*^ allele in some germ cells. Analysis of Cas9 levels revealed that transgene expression was much higher in testes than ovaries for both *Vasa-Cas9* lines, in contrast to the published Vasa-Cre line (Gallardo *et al*. 2007). Consistent with the sexually dimorphic expression level of each line, indel generation was higher in males than females. However, it remains unclear why indels were generated much more frequently in the *Vasa-Cas9-4* line versus *Vasa-Cas9-2* given that transgene expression in the testes was similar.

Recent studies indicate the level and timing of Cas9 expression in the germline is critical for homing efficiency. It is thought that homing is most likely to occur during meiosis when homologous chromosomes are aligned for meiotic recombination (Grunwald *et al*. 2019). Generation of DSBs before or after meiosis instead promotes generation of indels via error-prone repair pathways. As the timing of meiosis is sex-specific, activity of the Vasa transgene must be considered in both males and females. It has previously been shown that the Vasa promoter fragment is strongly induced at E15-18 in testes and before P3 in ovaries (Gallardo *et al*. 2007). In the E15-18 testes, primordial germ cells (PGCs) are undergoing mitotic proliferation and don’t enter meiosis until ∼P9 (Hilz *et al*. 2016). As a consequence, Vasa expression is likely too early for homing, consistent with the high frequency of indels that we observed in both Vasa-Cas9 lines. In contrast, nascent oocytes enter meiosis at E13.5, moving through zygotene to pachytene before they arrest at ∼E19 in dictyotene, which is maintained until ∼P21 (Hilz *et al*. 2016). As alignment of homologous chromosomes is maintained throughout this period, homing should be promoted. Why then did we not observe Super-Mendelian transmission of the *Tyr*^*gRNA*^ alleles in females? The answer probably relates to the very low level of transgene expression in the ovary, which, despite the substantial transgene copy number, was barely above background levels. Given only 9% and 11% of female *Vasa-Cas9* offspring carried indels, the frequency of DSB generation in the female germline was likely very low and homing rarely occurred if at all. It is also possible that the Vasa promoter is activated too early in the ovary given that endogenous

Vasa expression starts at E10.5-11.5 in PGCs of both sexes (Tanaka *et al*. 2000). Notably, in contrast to our results, a similar study by Grunwald *et al*. (2019) did observe homing in females (although not in males). An important difference between our study and Grunwald *et al*. (2019) is their use of the Vasa-Cre line to express Cas9 from the CMV promoter via removal of a stop-flox cassette. Thus, Cas9 levels are likely much higher in the Grunwald *et* al. (2019) experiment.

In conclusion, based on our observations and those of Grunwald *et al*. (2019), we suggest that zygotic homing is not a feasible strategy in mice. Efficient homing in the germline will require identification of additional promoters, ideally with robust and specific expression during meiosis in oocytes and spermatocytes. Thus, considerable experimental development is required before rodent gene drives can be considered for deployment to address conservation, agricultural or health objectives.

## Acknowledgements

This study was funded by a US Defense Advanced Research Projects Agency (DARPA) “Safe genes” grant to Paul Thomas. The authors thank John Godwin and members of the Genetic Biocontrol for Invasive Rodents consortium for useful discussions during the course of this research project. The authors acknowledge the facilities and the scientific and technical assistance of the South Australian Genome Editing (SAGE) Facility, the University of Adelaide and the South Australian Health and Medical Research Institute. SAGE is supported by the Australian Phenomics Network (APN). The APN is supported by the Australian Government through the National Collaborative Research Infrastructure Strategy (NCRIS) program.

