## Supplemental Data for "Development of zygotic and germline gene drives in mice"

### Supplemental Material

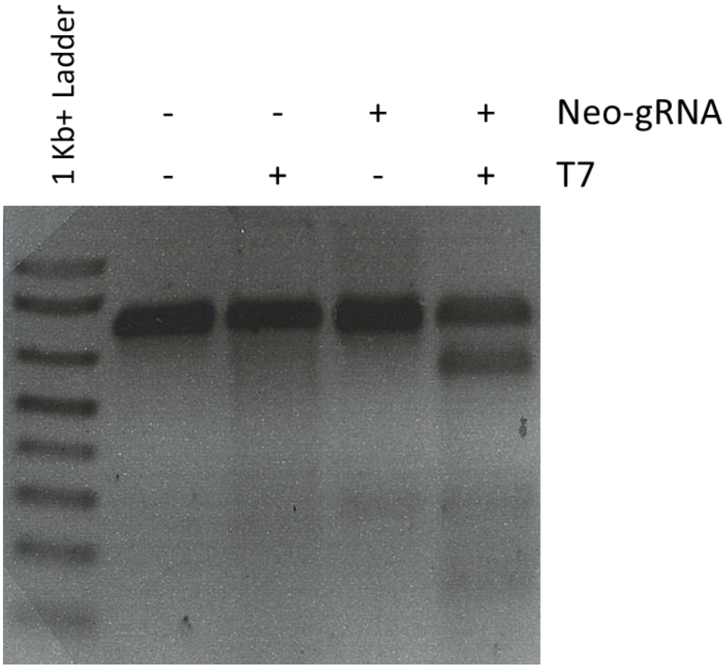


**Supplemental Figure 1. Demonstration of Neo-gRNA cleavage activity at its target binding site.** Mouse ES cells containing the target binding site for Neo-gRNA were transfected with a plasmid containing U6-driven Neo-gRNA and a plasmid (pX459) containing CMV-driven *hSpCas9*. RFLP analysis was performed around the cut site. Digestion of Neo-gRNA/T7 band demonstrates successful cutting by Neo-gRNA.

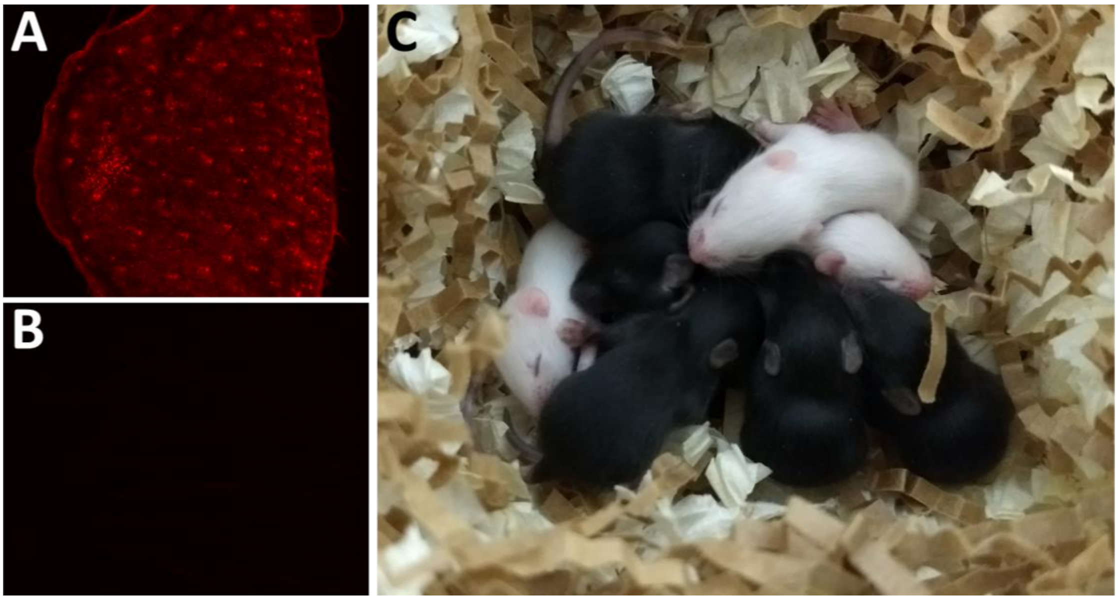


**Supplemental Figure 2. Phenotyping *Tyr*$^{gRNA\text{-}Tomato}$ mice. (A, B)** Representative immunofluorescence images of ear notches of **(A)** *Tyr*$^{gRNA\text{-}Tomato}$ mice and **(B)** WT mice, showing expression of dTomato. **(C)** A representative F2 litter of *Tyr*$^{gRNA\text{-}Tomato}$ mice. Black mice were heterozygous for *Tyr*$^{gRNA\text{-}Tomato}$. White mice were homozygous for *Tyr*$^{gRNA\text{-}Tomato}$.

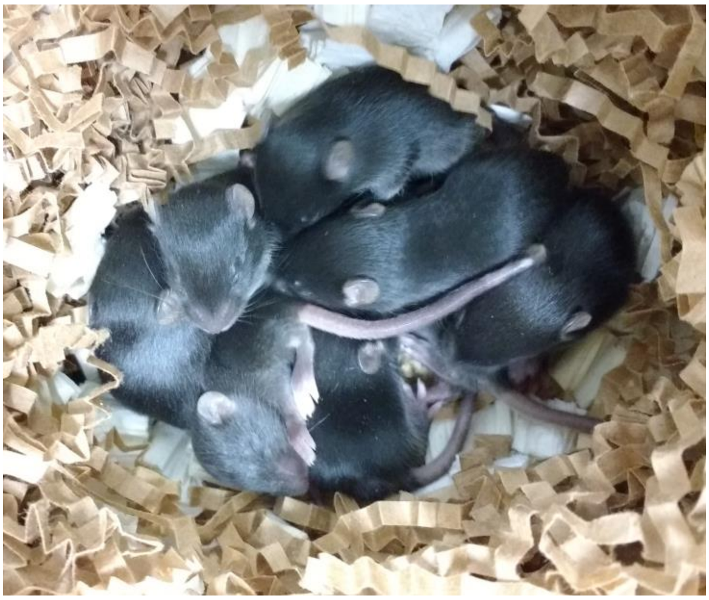

**Supplemental Figure 3.** *Tyr*$^{Target}$ homozygote phenotyping, showing black coats.

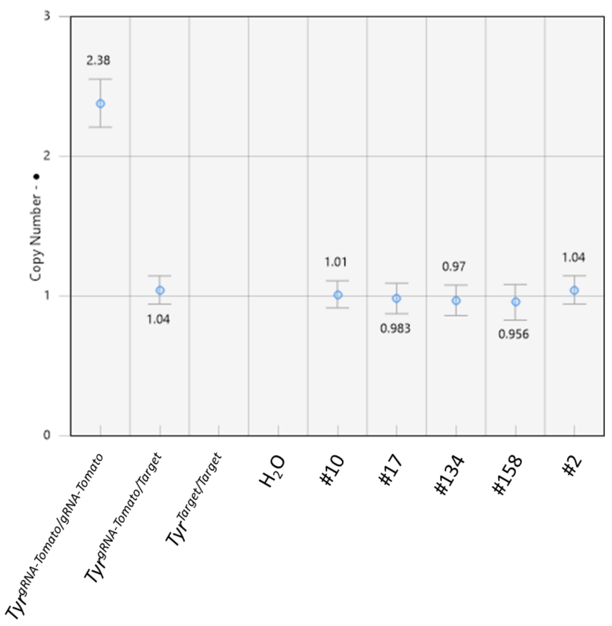


**Supplemental Figure 4. *Tyr*$^{gRNA\text{-}Tomato}$ copy number determination for zygotic-homing mice. Representative** ddPCR showing copy number of *Tyr*$^{gRNA\text{-}Tomato}$ (with 95% CI) in five *Rosa26*$^{Cas9}$/+ ; *Tyr*$^{gRNA\text{-}Tomato/Target}$ mice with dappled coats.

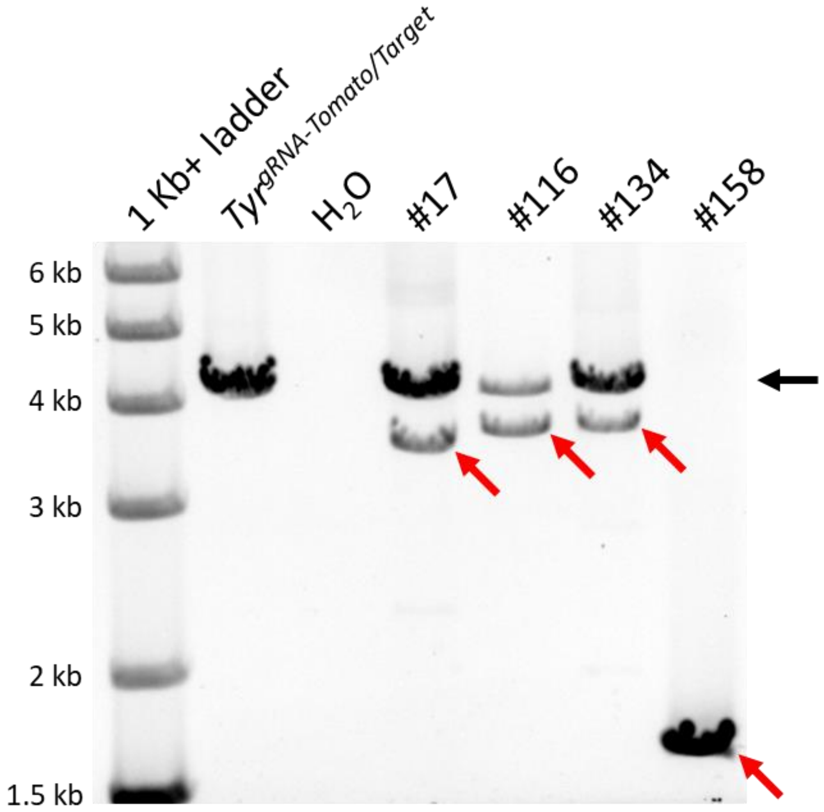


**Supplemental Figure 5. Large deletion genotyping.** PCR showing large deletions around $Tyr^{Target}$ in four $Rosa26^{Cas9}$/+ ; $Tyr^{gRNA\text{-}Tomato/Target}$ mice with dappled coats. The black arrow shows expected band (~4 kb) for no deletion/small indels. The red arrows show large deletions of varying size.

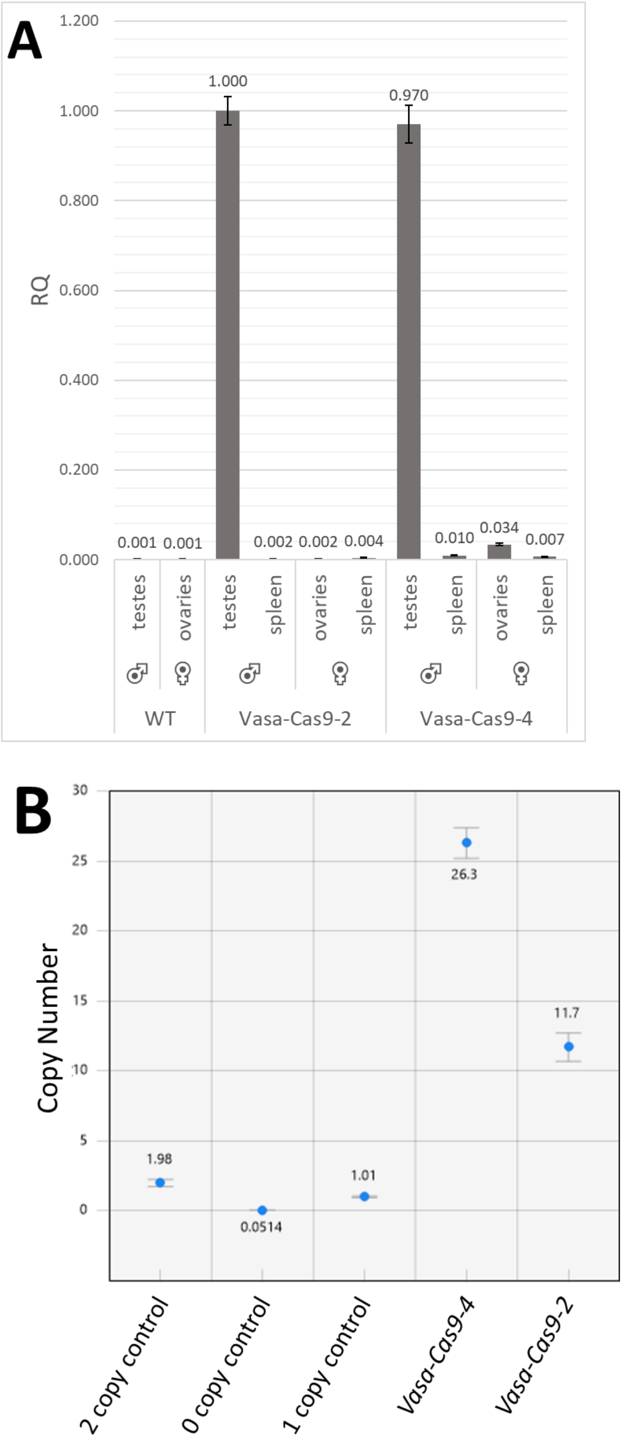


**Supplemental Figure 6. *Vasa-Cas9* expression. (A)** RT-qPCR showing expression levels (with 95% CI) of the *Vasa-Cas9* lines in various tissue types. **(B)** ddPCR copy number assay showing the copy number (with 95% CI) of the construct in *Vasa-Cas9-2* and *Vasa-Cas9-4* hemizygotes.

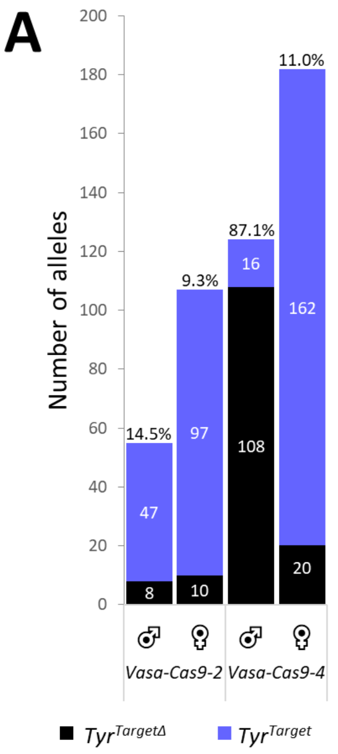


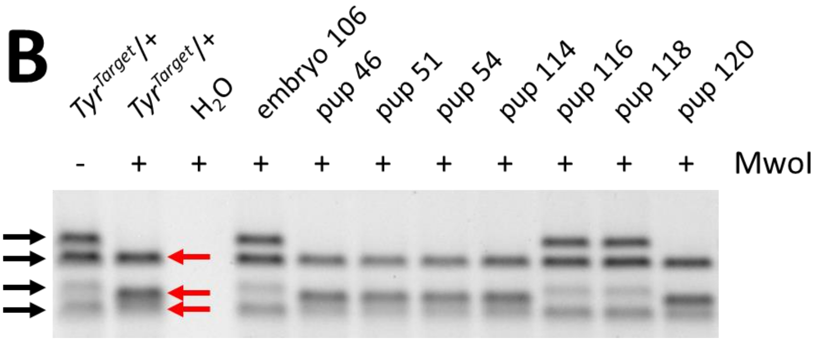


**Supplemental Figure 7. Extended germline-homing genotyping data. (A)** Data shows count of the *Tyr*<sup>*Target*</sup> alleles in the offspring of gene drive mice (as per Figure 3), broken down into *Vasa-Cas9* line and sex. *Tyr*<sup>*TargetΔ*</sup> alleles show the presence of an indel whereas *Tyr*<sup>*Target*</sup> alleles remain intact. Percentage of indels is shown at the top of each column. **(B)** Representative *Tyr*<sup>*Target*</sup> cut site digestion with MwoI (as indicated) and T7 Endonuclease (all samples), black arrows show expected uncut bands due to destruction of MwoI site indicating presence of an indel, red arrows show cut bands due to intact MwoI site and thus no indel.

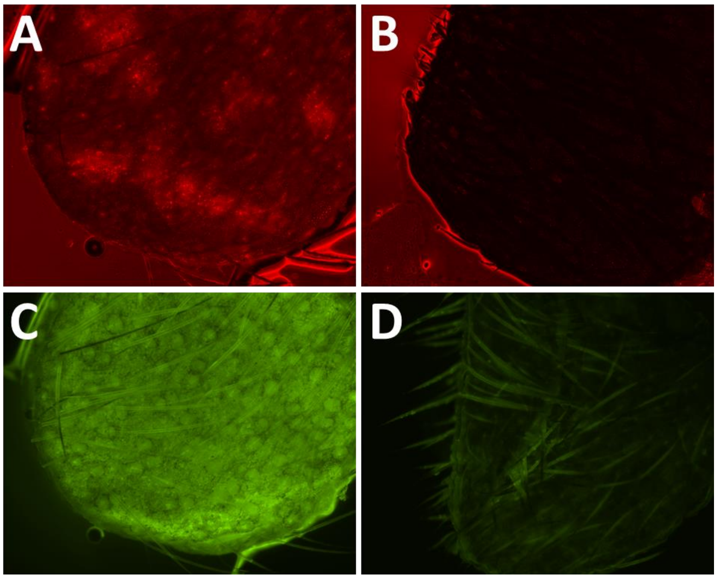


**Supplemental Figure 8. Phenotyping of *Rosa26*$^{Cas9}$/+ ; *Tyr*$^{gRNA\text{-}Tomato/Target}$ mice containing no indels.** Representative immunofluorescence images of ear notches of aforementioned mice **(A, C)** and WT mice **(B, D)**, showing expression of dTomato **(A, B)** and expression of EGFP **(C, D)**.

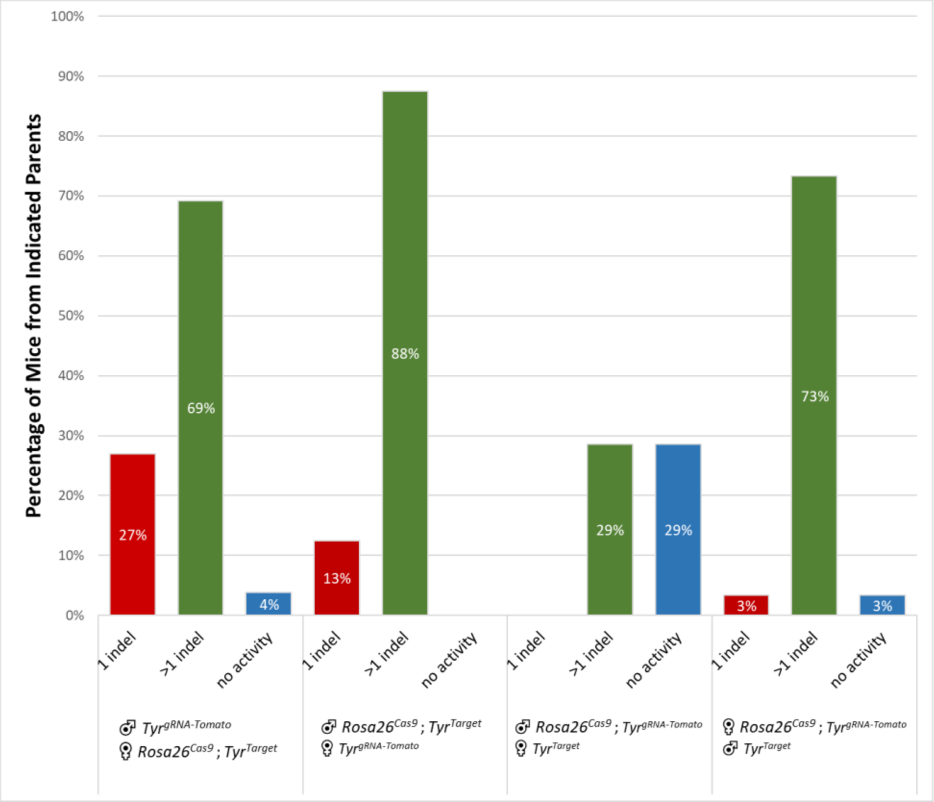


**Supplemental Figure 9. Breakdown of zygotic-homing indel formation based on parent of origin.** Sex symbols at bottom show which alleles were inherited from which parents.

| Mouse ID | Sex | *Tyr*$^{gRNA\text{-}Tomato}$ alleles | Total *Tyr*$^{Target}$ alleles | *Tyr*$^{Target\Delta}$ alleles |
|---|---|---|---|---|
| 3.2b | male | 23 | 18 | 4 |
| 27.1d | male | 16 | 17 | 3 |
| 27.1e | male | 3 | 6 | 0 |
| 33.2e | male | 2 | 5 | 0 |
| 33.2f | male | 9 | 3 | 0 |
| 33.2g | male | 11 | 6 | 1 |
| 24.1a | female | 8 | 5 | 0 |
| 33.2a | female | 19 | 14 | 0 |
| 33.2b | female | 3 | 5 | 0 |
| 35.1b | female | 3 | 1 | 1 |
| 35.1d | female | 4 | 4 | 0 |
| 35.1e | female | 3 | 3 | 0 |
| 35.2a | female | 24 | 27 | 3 |
| 35.2c | female | 0 | 8 | 4 |
| 36.2d | female | 21 | 26 | 1 |
| 37.2a | female | 8 | 6 | 1 |
| 37.2b | female | 10 | 3 | 0 |
| 37.2c | female | 12 | 19 | 0 |

**Supplemental Table 1. Germline-homing individual mouse genotyping data for *Vasa-Cas-2*.** Breakdown of the ratio of inherited alleles for the offspring of the listed *Vasa-Cas9-2*/+ ; *Tyr*$^{gRNA\text{-}Tomato/Target}$ mice crossed to WT. Data collated to produce *Vasa-Cas-2* data in **Figure 4** and **Supplemental Figure 7**.

| Mouse ID | Sex | *Tyr*$^{gRNA\text{-}Lite}$ alleles | Total *Tyr*$^{Target}$ alleles | *Tyr*$^{Target\Delta}$ alleles |
|---|---|---|---|---|
| 1.2e | male | 15 | 17 | 13 |
| 1.3f | male | 7 | 10 | 8 |
| 1.3g | male | 9 | 12 | 11 |
| 2.1f | male | 20 | 16 | 16 |
| 2.2d | male | 13 | 17 | 17 |
| 3.2g | male | 15 | 13 | 10 |
| 4.3f | male | 12 | 7 | 6 |
| 4.1g | male | 8 | 6 | 5 |
| 5.2d | male | 14 | 18 | 16 |
| 6.1e | male | 12 | 14 | 13 |
| 1.1b | female | 21 | 21 | 4 |
| 1.2b | female | 28 | 32 | 3 |
| 3.1b | female | 24 | 23 | 0 |
| 3.2e | female | 12 | 6 | 0 |
| 4.1b | female | 27 | 24 | 1 |
| 4.2b | female | 27 | 15 | 1 |
| 4.3d | female | 17 | 20 | 1 |
| 5.2c | female | 11 | 9 | 1 |
| 6.1b | female | 8 | 14 | 2 |
| 6.1c | female | 14 | 12 | 1 |

**Supplemental Table 2. Germline-homing individual mouse genotyping data for *Vasa-Cas-4*.** Breakdown of the ratio of inherited alleles for the offspring of the listed *Vasa-Cas9-4*/+ ; *Tyr*$^{gRNA\text{-}Tomato/Target}$ mice crossed to WT. Data collated to produce *Vasa-Cas-4* data in **Figure 4** and **Supplemental Figure 7**.

| Locus | Primer 1 | Primer 2 | Primer 3 | Type |
|---|---|---|---|---|
| *Tyr*$^{gRNA-Tomato}$ | CCAGACAGCCCTTGTAATCATTAGC | GGCTATCGTGGCGTTTTAGA | | PCR |
| *Tyr*$^{gRNA-Lite}$ | CCAGACAGCCCTTGTAATCATTAGC | AACTTGAAAAAGTGGCACCGAG | GCACCTCCTATGGTATCTGGAA | PCR |
| *Tyr*$^{Target}$ | ACTGTTTGAGAGTCAGCAACGT | TCTCTGGCCAAAACCAAGACTT | | PCR/RFLP |
| *Tyr*$^{Target}$ | GGGTTCTGTCCTCAACTGGT | TTTGATGTAAGAAGGGGAGTGGT | | Large Δ PCR |
| *Rosa26*$^{Cas9}$ | AAGGGAGCTGCAGTGGAGTA | CCGAAAATCTGTGGGAAGTC | CCATAAGGTCATGTACTGGGC | PCR |
| *Vasa-Cas9* | ATTGTACTTCAGCACAGTTTTAGAG | AGTCTCCGTCGTGGTCCTTA | | PCR |
| *Vasa-Cas9* | ACCTGAACCCCGACAACA | CTGGCGTTGATGGGGTTTTC | | RT-qPCR |

**Supplemental Table 3. List of PCR primers.**
